# DNA damage-associated vesicle production in *Stenotrophomonas maltophilia* is mediated by a cryptic tailocin endolysin

**DOI:** 10.1101/2025.03.20.644284

**Authors:** Darshan Chandramowli, Jolien Vitse, Stephan Stremersch, Koen Raemdonck, Kevin Braeckmans, Bart Devreese

## Abstract

Like other Gram-negative bacteria, *S. maltophilia* is capable of producing membrane vesicles under normal growth conditions. The addition of certain exogenous triggers stimulates the production of vesicles, including those that are distinct from the archetypal outer membrane vesicle. In this study, we examine the effects of DNA damage on vesiculation, with a focus on the role of a bacteriophage-encoded endolysin. We demonstrate that deletion of the gene that encodes this protein (*mal*) negatively affects the vesiculation capacity of *S. maltophilia*. Further, we provide evidence that the spontaneous re-arrangement of shattered membrane fragments attributable to *mal*-induced explosive cell lysis results in the formation of predominantly explosive outer membrane vesicles, while only a minor portion are outer-inner membrane vesicles. Our findings expand on the current knowledge of (cryptic) tailocins in the biogenesis of vesicles in conditions of stress.

## Introduction

Since the first observation of membrane vesicles (MVs) in a culture of *E. coli* in the mid-1960s, several reports of MVs produced by other organisms have been published, leading to the hypothesis that vesiculation is a conserved process across all domains of life ^1,2^. In the decades since, our knowledge about the different types of MVs and their functions has signficantly improved – despite this, it has also been posited that our current perceptions severely underestimate their roles ^3,4^. In Gram-negative bacteria, the most common type of MVs produced result from the blebbing of the outer membrane, giving rise to outer membrane vesicles (OMVs). Various triggers can result in the production of OMVs, including oxidative stress and exposure to cell wall-destabilising antibiotics like β-lactams ^3^. Interestingly, the cargo enveloped within OMVs has been shown to correlate with the trigger that stimulated its production – for example, OMVs produced by the opportunistic pathogen *Stenotrophomonas maltophilia* exposed to a sub-minimum inhibitory concentration (MIC) of benzylpenicillin were found to be enriched with two chromosomally-encoded β-lactamases (L1 and L2), and administration of these vesicles to cultures of *Pseudomonas aeruginosa* and *Burkholderia cenocepacia* was demonstrated to protect against imipenem and ticarcillin ^5,6^.

One surprising observation from early experiments was the detection of genetic material packaged into MVs ^7–9^. The presence of these cytoplasmic contents could not be explained under the assumption that vesicles were produced exclusively from the outer membrane – as a result, it was theorised that these represented a new type of MV formed by the protrusion of the inner cell membrane into the periplasm, such that the resulting vesicle would have both outer and inner membranes. The first experimental evidence for this type of vesicle came from experiments conducted on *Shewanella vesiculosa*, where transmission electron microscope (TEM) images unequivocally revealed the presence of vesicles with a bilayered membrane – these were aptly called outer-inner membrane vesicles (O-IMVs or OIMVs) ^10^. Such vesicles have since been identified in multiple other pathogenic bacteria, including *P. aeruginosa* and *Acinetobacter baumannii* ^11^.

When faced with DNA damage, one of the first responses in bacteria is the upregulation of the SOS response. This system is mediated by two proteins – the recombination protein RecA and the transcriptional repressor LexA. Briefly, RecA binds to ssDNA resulting from DNA damage to form a nucleoprotein filament which interacts with LexA and causes its autocatalytic cleavage. This allows LexA-repressed genes (called SOS genes) to be upregulated and repair the damaged DNA through rapid (but error-prone) pathways ^12^. One important consequence of this global response is the upregulation of lysogenic bacteriophages, such as the λ phage in *E. coli* ^13^. As a consequence, they enter the lytic stage of their life cycles, wherein the host bacterial cells are lysed to release multiple progeny viruses that can infect other cells. Recently, it was proposed that such an event (termed explosive cell lysis) was responsible for the production of OIMVs in *P. aeruginosa*. It was demonstrated that upregulation of the SOS response stimulated the production of a prophage endolysin (*lys*) which acts on the peptidoglycan layer of the cell wall, weakening it enough to cause cells to ‘explode,’ with the spontaneous rearrangement of membrane fragments into OIMVs as a consequence ^14^. This pathway is now considered to be the major mechanism of OIMV biogenesis, although this is still debated by some as a ‘true’ biogenesis route since it typically involves cell death ^3^.

Perhaps unsurprisingly, OIMVs have also been detected in culture supernatants of *S. maltophilia* exposed to ciprofloxacin ^15^. Fluoroquinolones (such as ciprofloxacin) function by inhibiting the actions of bacterial DNA gyrase and topoisomerases, resulting in single and double-strand breaks in replicating DNA strands and subsequent cell death ^16^. Akin to the more well-characterised *P. aeruginosa*, there exists a cryptic prophage within the genome of *S. maltophilia*, first described in the strain P28. This tailocin (named maltocin) is encoded by a gene cluster that shows similarity to the organisation of the rigid (R)-type pyocin gene cluster in *P. aeruginosa* and has antibacterial activity against other strains of *S. maltophilia* ^17^. One gene in this maltocin operon (*smlt1054* in the clinical type strain K279a) is predicted to code for an endolysin that shows broad spectrum bactericidal activity ^18^. In this study, we work with the hypothesis that this endolysin (which we designate LysSM in this study) plays an analogous role to the *lys* endolysin in *P. aeruginosa* for the production of MVs in response to DNA damage. We report on the effects of antibiotic and genotoxic-mediated DNA damage stress on the membrane integrity of a clinical isolate of *S. maltophilia* (strain 44/98, LMG 26824) in LysSM deletion mutant (Δ*mal*) using microscopy and other microbiological techniques.

## Materials and methods

### Bacterial strains and growth conditions

Strains and plasmids used in this study are listed in Table 1. *E. coli* was maintained in regular LB (LB-Miller), while *S. maltophilia* was maintained in a low-salt LB formulation (LB-Lennox). *S. maltophilia* was also grown on *Pseudomonas* Isolation Agar (PIA) plates for selection of successful transformants. All liquid cultures were grown at 37°C in aerobic conditions, and agar plates were incubated at either 30 or 37°C. The following antibiotics were used for selection in or against *E. coli* – trimethoprim at 50 μg/ml, kanamycin at 50 μg/ml, tetracycline at 10 μg/ml, and norfloxacin at 2 μg/ml. The following antibiotics were used for selection in *S. maltophilia* – trimethoprim at 50 μg/ml, chloramphenicol at 40 μg/ml, and tetracycline at 15–17 μg/ml. The following antibiotics were used for induction of stress responses in *S. maltophilia* – imipenem at 25 μg/ml, carbenicillin at 200 μg/ml, ciprofloxacin at 2 μg/ml, norfloxacin at 2 μg/ml, and mitomycin C at 1 μg/ml.

**Table 1.**
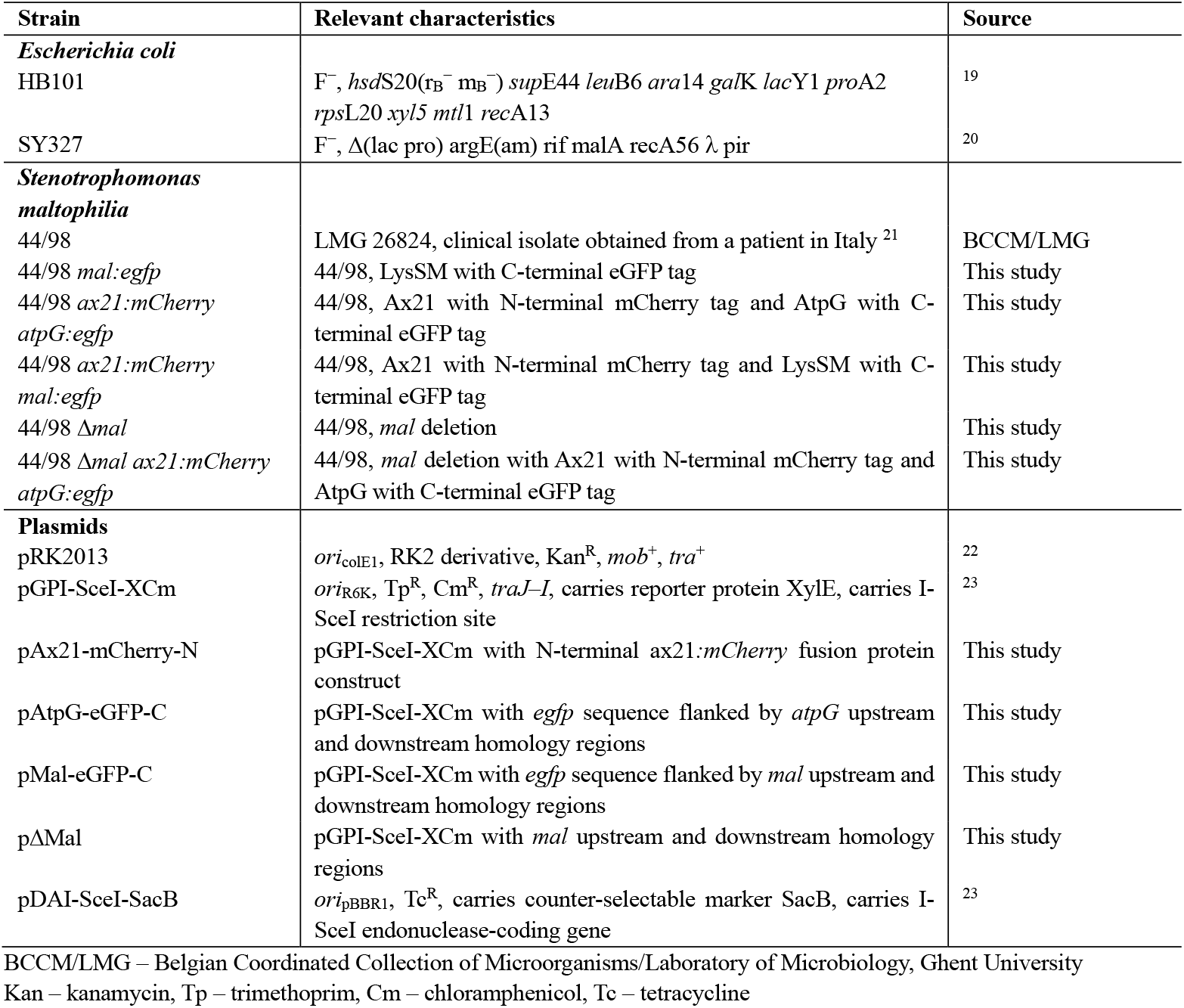
Strains and plasmids used in this study.

### Construction of *ax21:mCherry, atpG:egfp, mal:egfp*, and Δ*mal* mutants

The 44/98 deletion and insertion mutants used in this study were constructed using the protocol described previously ^24^. The *ax21:mCherry* mutants were constructed by replacing the endogenous *ax21* gene with the corresponding mCherry-tagged version, while the *atpG:egfp* and *mal:egfp* mutants were constructed as in-frame fusions with *egfp*. Construction of all plasmids was outsourced to GenScript Biotech (Piscataway, New Jersey). Verification of all mutations was performed by sequencing of PCR fragments generated using various primer pairs (Supplementary Material 1).

To construct the *ax21:mCherry* replacement allele, regions upstream (393 bp) and downstream (500 bp) of *ax21* were selected. The upstream region was comprised of mostly intergenic sequences and a small portion of the preceding gene (*smlt0386*), while the downstream region was made up of entirely intergenic sequence. The replacement gene itself was comprised of the mCherry-coding sequence placed *after* the signal peptide sequence, followed by a short flexible linker (coding for the amino acid sequence GSGSGS) and the gene itself (Supplementary Material 2). The entire allele (2.5 kb) was synthesised and cloned into pGPI-SceI-XCm to give pAx21-mCherry-N.

To construct the insert allele for *atpG*, regions upstream (839 bp) and downstream (861 bp) of the gene were selected. The upstream region consisted of a portion of the gene immediately upstream (*atpD*), while the downstream region was comprised of the gene itself *without* its stop codon. Since *atpG* is found on the minus-strand, the reverse complements of the eGFP-coding sequence and linker were used. The final insert was made up of the upstream homology sequence (*atpD*), the eGFP and linker (coding for the amino acids GSGSGS) sequences, and the downstream homology sequence (*atpG*) (Supplementary Material 2). This sequence was synthesised and cloned into pGPI-SceI-XCm to give pAtpG-eGFP-C.

To construct the insert allele for *mal*, regions upstream (200 bp) and downstream (300 bp) of the gene were selected. The upstream region consisted of a portion of the gene immediately upstream (*smlt1053*), while the downstream region was comprised of a portion of the gene itself *without* its stop codon. *mal* is also found on the minus-strand, hence the reverse complements of the eGFP-coding sequence and linker (coding for the amino acids GSGSGS) were used once again. The final insert was made up of the upstream homology sequence (*smlt1053*), the eGFP and linker sequences, and the downstream homology sequence (*smlt1055*) (Supplementary Material 2). This sequence was synthesised and cloned into pGPI-SceI-XCm to give pMal-eGFP-C.

To construct the deletion allele for *mal*, regions upstream (200 bp) and downstream (300 bp) of the gene were selected. The upstream portion consisted of a portion of the gene immediately upstream (*smlt1053*), while the downstream region was comprised of a portion of the gene immediately downstream (*smlt1055*) (Supplementary Material 2). This sequence was synthesised and cloned into pGPI-SceI-XCm to give pΔ Mal.

### Growth curve construction and specific growth rate calculation

Overnight cultures of *S. maltophilia* 44/98 wild-type (WT) and Δ *mal* were back-diluted to an OD_600_ of 0.01 in 50 ml of LB-Lennox. Cultures were incubated in a rotary shaker at 37°C and 200 rpm, and OD_600_ measurements were taken at intervals of 30 minutes for 12 hours. In addition to monitoring the growth of the WT and Δ *mal* mutant under normal conditions, growth curves were also constructed for cultures exposed to sub-MIC concentrations of norfloxacin and mitomycin C in parallel – for these experiments, the corresponding antibiotic was added once cultures had reached an OD_600_ between 0.3 and 0.4. Specific growth rates were calculated for each growth condition between 4 and 10 hours (corresponding to the exponential phase) from the graphs plotted as previously described ^25^. Growth curves were constructed using three biological replicates per condition.

### Biofilm assay

Overnight cultures of *S. maltophilia* 44/98 WT and Δ *mal* were normalised to an OD_600_ of 2.0 and subsequently back-diluted in a ratio of 1:100 in a 3-(*N*-morpholino)propanesulphonic acid (MOPS) minimal medium supplemented with 5 mM glucose and 250 μM L-methionine ^26^. 100 μl of these suspensions were inoculated in each well of separate 96-well flat-bottom polystyrene plates. Plates were sealed using microporous sheets (Qiagen) and incubated statically at 37°C for 4 hours to allow cell to adhere. Following this, the medium was discarded and cells were washed with 150 μl of sterile phosphate-buffered saline (PBS) to remove non-adherent cells. 100 μl of fresh MOPS medium was added to each well and plates were sealed once again. Three conditions of stress were tested per plate – MOPS without any antibiotic, and MOPS supplemented with sub-MICs of either norfloxacin or mitomycin C. Three biological conditions with eight technical replicates each were tested per condition. Plates were incubated statically at 37°C for 24 or 48 hours to allow biofilm formation.

After overnight incubation, the medium was discarded and cells were washed with 150 μl of sterile PBS to remove non-adherent cells. Cells were fixed with 99 % (v/v) methanol for 15 minutes, following which methanol was aspirated and plates were air-dried. 100 μl of 0.01 % (w/v) crystal violet was added to each well and plates were incubated for 30 minutes at ambient temperature. Following staining, plates were washed thoroughly under running tap water to remove unbound crystal violet. Bound crystal violet was liberated by addition of 150 μl of 33 % (v/v) glacial acetic acid and absorbance was measured at 590 nm. Prior to plotting graphs, absorbance values were normalised to specific growth rates for each condition to account for differences in growth. Unpaired t-tests were performed using an online t-test calculator (GraphPad) to assess the significance of results.

### Isolation of MVs

Overnight cultures of *S. maltophilia* 44/98 WT and Δ*mal* cultures were back-diluted in 50 ml of fresh LB-Lennox at a ratio of 1:50 and allowed to grow until they reached mid-exponential phase (OD_600_ between 0.65–0.75). Vesiculation was stimulated by addition of a sub-MIC concentration of β-lactam (imipenem or carbenicillin), fluroquinolone (ciprofloxacin or norfloxacin), or mitomycin C, and cultures were allowed to grow for 3 hours. Subsequently, cultures were centrifuged at 8000 x g for 10 minutes at 4°C to pellet cells, and the supernatant was passed through a 0.45 μm pore-size polyethersulphone membrane filter (Merck Millipore, Massachussets). 40 ml of filtered supernatant was transferred to a polycarbonate vial and ultra-centrifuged at 108800 x g for 3 hours at 4°C (Avanti J-30I, Beckman Coulter, California) to pellet MVs. The MV pellet was resuspended in phosphate-buffered saline (PBS) and stored at –20°C until required.

### MV quantification and size determination

The concentration and diameters of MVs were determined by single particle-tracking (SPT) using a NanoSight LM10 HS system (Malvern Panalytical, United Kingdom) equipped with a 405 nm laser. Movies of 30 seconds were recorded and analysed using the NTA Analytical Software version 2.3 (Malvern Panalytical, United Kingdom). Calculations were made as previously described ^27^. Unpaired t-tests were performed using an online t-test calculator (GraphPad) to assess the significance of results.

### Fluorescence microscopy

Overnight cultures of *S. maltophilia* mutants with fluorophore insertions were back-diluted in fresh LB-Lennox at a ratio of 1:50 and allowed to grow for 3 hours. When the cultures reached an OD_600_ between 0.3 and 0.4, the appropriate antibiotic was added, and cultures were grown for an additional 2 hours. Meanwhile, plastic microscopy dishes (refractive index 1.5) were coated with 0.1 % (w/v) poly-L-lysine for 20 minutes, following which they were washed with sterile MilliQ water and air-dried in a laminar flow hood. For imaging, 200 μl of each culture was pipetted on to each poly-L-lysine-coated dish and placed in a laminar flow hood for 20 minutes. After incubation, the excess culture was removed and plates were allowed to air-dry for an additional 10 minutes. A drop of immersion oil (refractive index 1.52) was placed on the dish before transferring it to the microscope stage. For time-lapse microscopy, samples were prepared in the same way described above – prior to imaging, 20 μl of high-concentration norfloxacin was added directly to air-dried plates for a duration of 1 minute and subsequently removed.

Samples were visualised using a spinning disk confocal microscope (Nikon Eclipse Ti, Japan) equipped with an MLC 400 B laser box (Agilent Technologies, California, USA), a Yokogawa CSU-X confocal spinning disk device (Andor, Belfast, UK), an iXon Ultra EMCCD camera (Andor Technology, Belfast, UK), and a Plan Apo VC 100× 1.4 NA oil immersion objective lens (MRD01902, Nikon, Japan). A 1.5× zoom lens was used for additional magnification on the camera. The NIS Elements software package (Nikon, Japan) was used for imaging. The 488 nm and 561 nm diode laser lines were applied sequentially to excite the eGFP and mCherry-tagged proteins respectively, and fluorescence emission was detected through a quad band filter (440/40, 521/21, 607/34, 700/45). Exposure time was set to 300 ms for mCherry and 1 s for eGFP, and 8× averaging was used to improve the signal-to-noise ratio. 16-bit images were recorded with an EM gain of 300 (without binning). The image size was 512 × 512 pixels, with a pixel size of 90 nm. For time-lapse videos, images were taken at 5 second intervals for a duration of 5 minutes each with the Perfect Focus System (PFS) enabled. Samples were maintained at ambient temperature during imaging. Images were analysed using Fiji ^28^.

### Transmission electron microscopy

400-mesh copper grids (Electron Microscopy Sciences, Pennsylvania) were coated with 3 nm carbon and glow-discharged for 40 seconds at 15 mA prior to loading samples. Samples were blotted on the coated sides of the grids, which were held between self-locking tweezers. After 1 minute, grids were carefully dried with filter paper and washed with 5 consecutive drops of distilled water. Grids were then carefully placed on top of a droplet of 25 % (w/v) uranyl acetate replacement stain (Electron Microscopy Sciences, Pennsylvania) for 1 minute. Excess stain was removed with filter paper, and grids were air-dried for at least 1 hour before imaging. Imaging was performed at 80 kV on a JEM-1400 Plus electron microscope (JEOL, Tokyo).

## Results

### Construction of Δ*mal* mutants

The Δ*mal* mutants were constructed using an allelic-exchange mutagenesis strategy optimised for poorly-characterised multidrug-resistant bacteria ^24^. Of note, we noted success when using homology regions signifcantly shorter (200–300 bp) than those typically reported in literature thus far (500–1500 bp) with comparable efficiency (Figure 1) ^29–31^. This finding stimulates future studies using this method to expend still less time by reducing the length of homology regions when constructing suicide plasmids, which is by far the most time-consuming and tedious part of the protocol.

**Figure 1.**
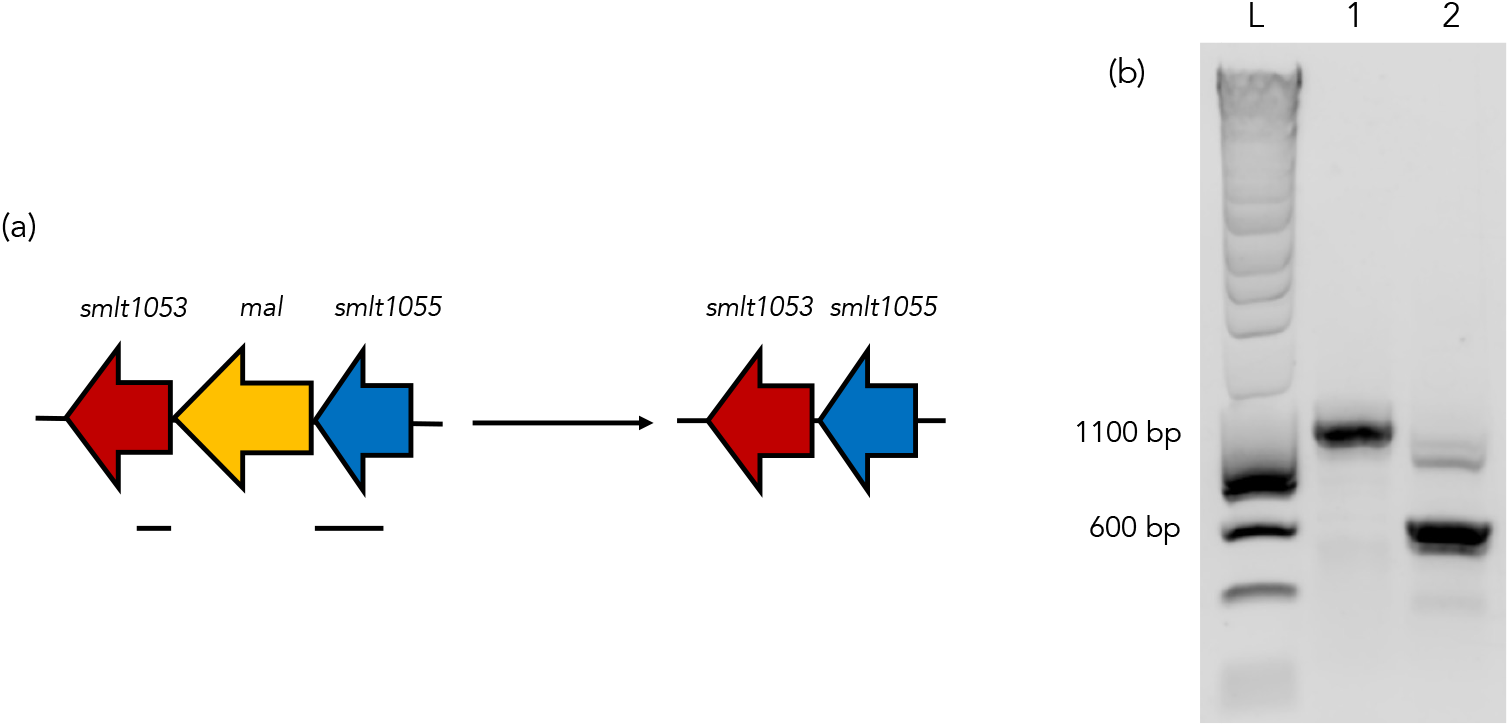
Deletion of *mal*. a) Deletion schematic for Δ*mal*. The upstream and downstream homology regions used for construction of the suicide plasmid are marked under the figure. b) PCR confirmation of Δ*mal*. L – ladder; 1 – PCR of WT flanking regions of *mal*; 2 – PCR of Δ*mal* flanking regions.

### Deletion of *mal* appears to have a slightly positive impact on *S. maltophilia* growth but impairs biofilm formation and drastically affects vesiculation

To investigate the effect of the deletion of *mal* on viability of *S. maltophilia*, growth curves were constructed for the WT and Δ*mal* strains. We observed that the deletion resulted in slightly positive growth under normal conditions of growth, so growth curves were constructed for both strains upon exposure to sub-MICs of norfloxacin and mitomycin C at the start of the exponential phase. Here too, it was observed that growth of the Δ*mal* mutant exposed to the DNA-damaging compounds was better than the corresponding growth in the WT (Figure 2). We rationalise these findings by correlating the absence of LysSM with decreased cell lysis relative to the WT, thereby improving cell growth.

**Figure 2.**
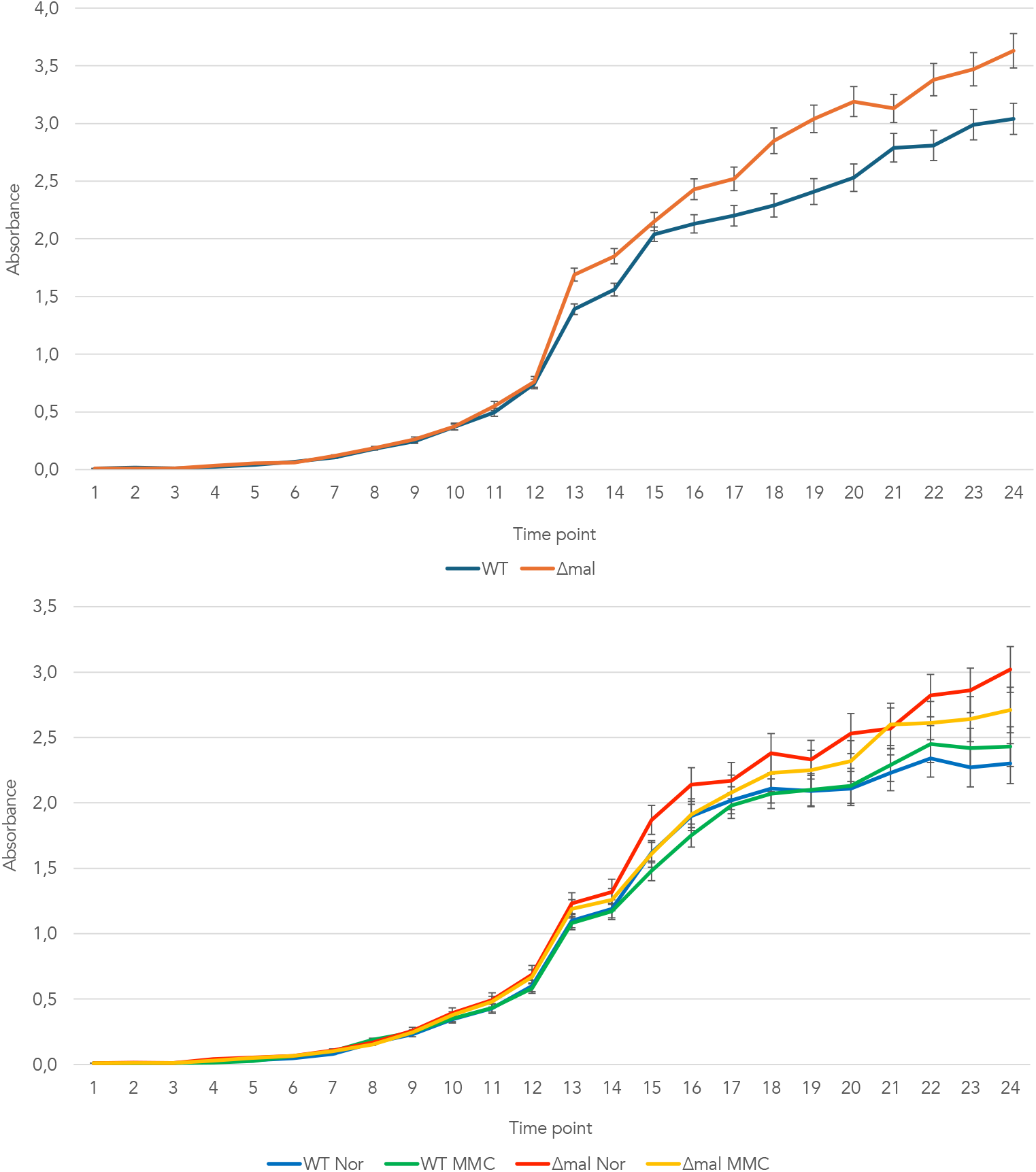
Growth curves of 44/98 WT and Δ*mal*. (above) Growth curves were constructed for cultures to which no antibiotics were added. (below) Growth curves were constructed for cultures to which sub-MICs of norfloxacin or mitomycin C were added. Time points shown on the x-axis correspond to intervals of 30 minutes. Nor – norfloxacin, MMC – mitomycin C.

To further test this hypothesis, we examined the effect of deletion of LysSM on biofilm forming capability and MV production. Cell lysis results in the release of extracellular DNA and various cytoplasmic contents, which are known to stimulate biofilm formation, and production of different types of MVs, most notably outer-inner membrane vesicles (OIMVs) and explosive outer membrane vesicles (E-OMVs). Concurrent with our hypothesis, we observed a decrease in biofilm formation after 24 and 48 hours in the Δ*mal* mutant relative to the WT when no antibiotics were added to the growth medium, as well as in response to DNA damaging agents (Figure 3a). Further, a significant decrease in MV formation was also observed between WT (1.16 × 10^11^ particles/ml) and Δ*mal* cultures (2.91 × 10^9^ particles/ml) (Figure 3b). Taken altogether, these findings highlight the role of LysSM in endolysin-triggered cell death and consequent MV production (Table 2).

**Table 2.** Statistical analysis of the effect of deletion of LysSM.

| Condition | p-value | Significance |
| --- | --- | --- |
| **Biofilm formation** |  |  |
| No antibiotic, 24 hours | 0.0338 | * |
| No antibiotic, 48 hours | 0.0096 | ** |
| Norfloxacin, 24 hours | 0.0362 | * |
| Norfloxacin, 48 hours | 0.0610 | n.s. |
| Mitomycin C, 24 hours | 0.0037 | ** |
| Mitomycin C, 48 hours | 0.0347 | * |
| **MV production** |  |  |
| WT vs. $\Delta mal$ | 0.0007 | *** |
| **MV size** |  |  |
| WT vs. $\Delta mal$ | 0.0001 | **** |

$p > 0.05$ n.s.; $p < 0.05$ \*; $p < 0.01$ \*\*; $p < 0.001$ \*\*\*; $p < 0.0001$ \*\*\*\*.

**Figure 3.**
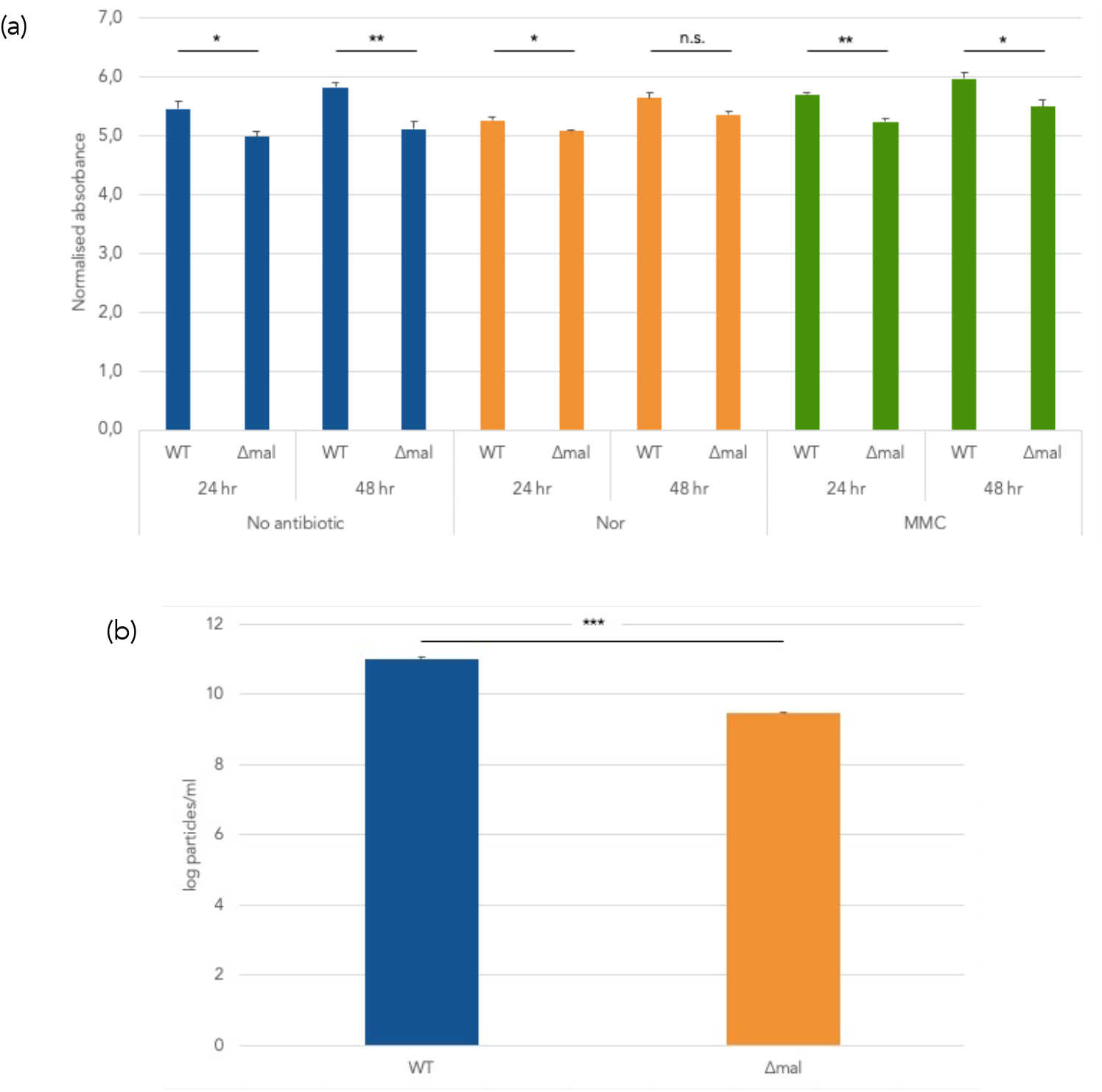
Effect of deletion of LysSM. a) Comparison of biofilm formation between 44/98 WT and Δ*mal* in conditions of no antibiotic exposure, exposure to norfloxacin, and exposure to mitomycin C. Graph is shown as mean ± S.E.M. b) Comparison of MV production between 44/98 WT and Δ*mal*. Plotted values have been log-transformed for the sake of convenience. Graph is shown as mean ± S.D.

Based on the results of the SPT experiment, a size distribution graph was constructed for the vesicles isolated from the WT and Δ*mal* cultures after exposure to ciprofloxacin. The optimal size range of the instrument used is around 100–1000 nm, so particles smaller than the lower limit are underrepresented. Based on the data, it was observed that the average size of MVs obtained from the WT cultures (267.7 nm) was significantly larger than those retrieved from the Δ*mal* cultures (169.24 nm) – further, the size range of WT vesicles was far greater than that of the Δ*mal* vesicles (Figure 4, Table 2). For both strains, the size of the resulting MVs were considerably larger than those previously shown to be formed in response to β-lactams like imipenem ^15^. This suggests that LysSM is involved in the production of MVs in *S. maltophilia* in response to DNA damage, and the MVs produced by this route are likely morphologically distinct from those produced in response to cell wall stress.

**Figure 4.**
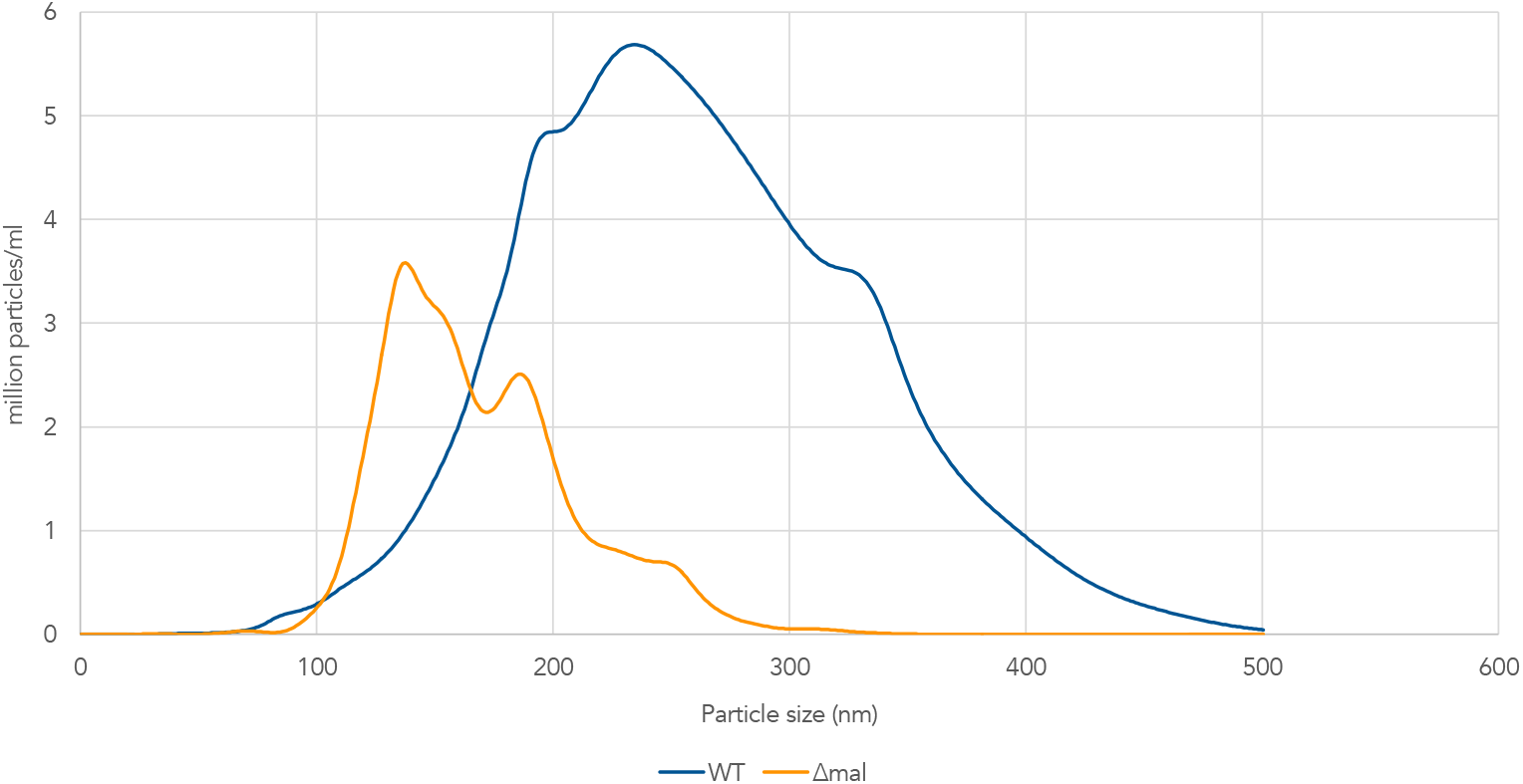
Size distribution of MVs produced by 44/98 WT and Δ*mal*. The average size of MVs produced by the WT is much larger than that of MVs isolated from the Δ*mal* mutant. Further, the size range of WT vesicles (around 100–500 nm) is much more pronounced than Δ*mal* vesicles (around 100–300 nm).

### LysSM is implicated in cell lysis and consequent MV production – however, E-OMVs are the predominant type of MV formed, not OIMVs

In order to study the discrepancy in average MV size between the WT and Δ*mal* mutant, fluorophore-tagged mutants were constructed to try and visualise the effect of DNA stress on membrane integrity. For this purpose, fluorophore-tagged mutants were constructed for the WT and Δ*mal* mutant, wherein the outer membrane protein Ax21 was tagged with mCherry and the inner membrane ATP synthase Ρ subunit AtpG was tagged with eGFP – the resulting strains are referred to as DT (i.e., double-tagged) and DTΔ respectively in the subsequent sections.

As an initial test, exponentially-dividing cultures of DT and DTΔ were subjected to carbenicillin for 2 hours. Exposure to such a cell wall-targeting antibiotic was anticipated to result in the production of exclusively OMVs – as expected, fluorescence microscopy revealed the presence of spherical particles that showed signals from the mCherry but not the eGFP channel in both samples (Figure 5). Curiously, it was observed that AtpG became localised at specific well-defined foci throughout cells in treated samples compared to a more uniform distribution in the corresponding untreated controls – to a lesser extent, Ax21 also appeared to concentrate at the polar regions of many cells (Figure 5, Supplementary Material 3). Spatiotemporal changes in protein localisation are considered good indicators of stress in cells; therefore, we were confident that our mutants could be used to assess membrane integrity ^32^.

**Figure 5.**
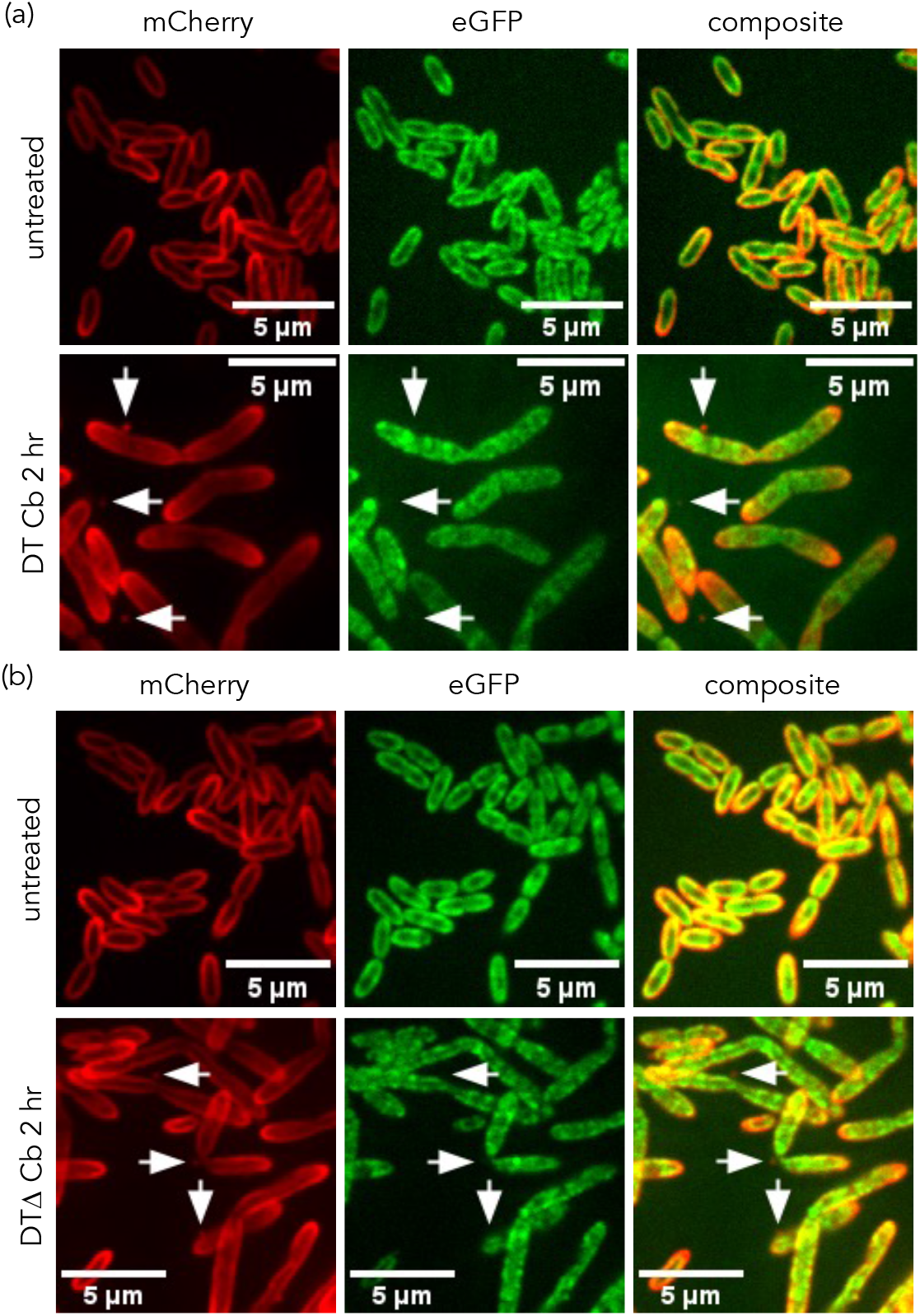
Effect of cell wall-targeting compounds on membrane integrity. (a) DT control and carbenicillin-treated cultures. Arrows show OMVs with mCherry signals visible and corresponding eGFP signals absent. (b) DTΔ control and carbenicillin-treated cultures. Arrows show OMVs with mCherry signals visible and corresponding eGFP signals absent. Re-localisation of Ax21 (outer membrane) and AtpG (inner membrane) are visible. Cb – carbenicillin.

We then sought to gauge the severity of the effect of DNA stress by exposing DT and DTΔΔ cultures to sub-MICs of either norfloxacin or mitomycin C overnight. After 24 hours, fluorescence microscopy revealed several bright spots separate from cells and distinct from background fluorescence, corresponding to both outer and inner membrane fragments. Analysis of the samples revealed two crucial observations – it was observed that spots were more abundant in DT samples when compared to DT?Δ samples, and overlapping of mCherry and eGFP signals was only accurately detected in DT samples (Figure 6, Supplementary Material 3). Similar results were seen when the experiment was repeated with only 2 hours of exposure to norfloxacin or mitomycin C (Figure 7). We conclude that the spots with coinciding fluorescence signals correspond to bilayered OIMVs formed by spontaneous re-circularisation of membrane fragments after LysSM-induced cell lysis, thus explaining why they were not often detected in DTΔ samples.

**Figure 6.**
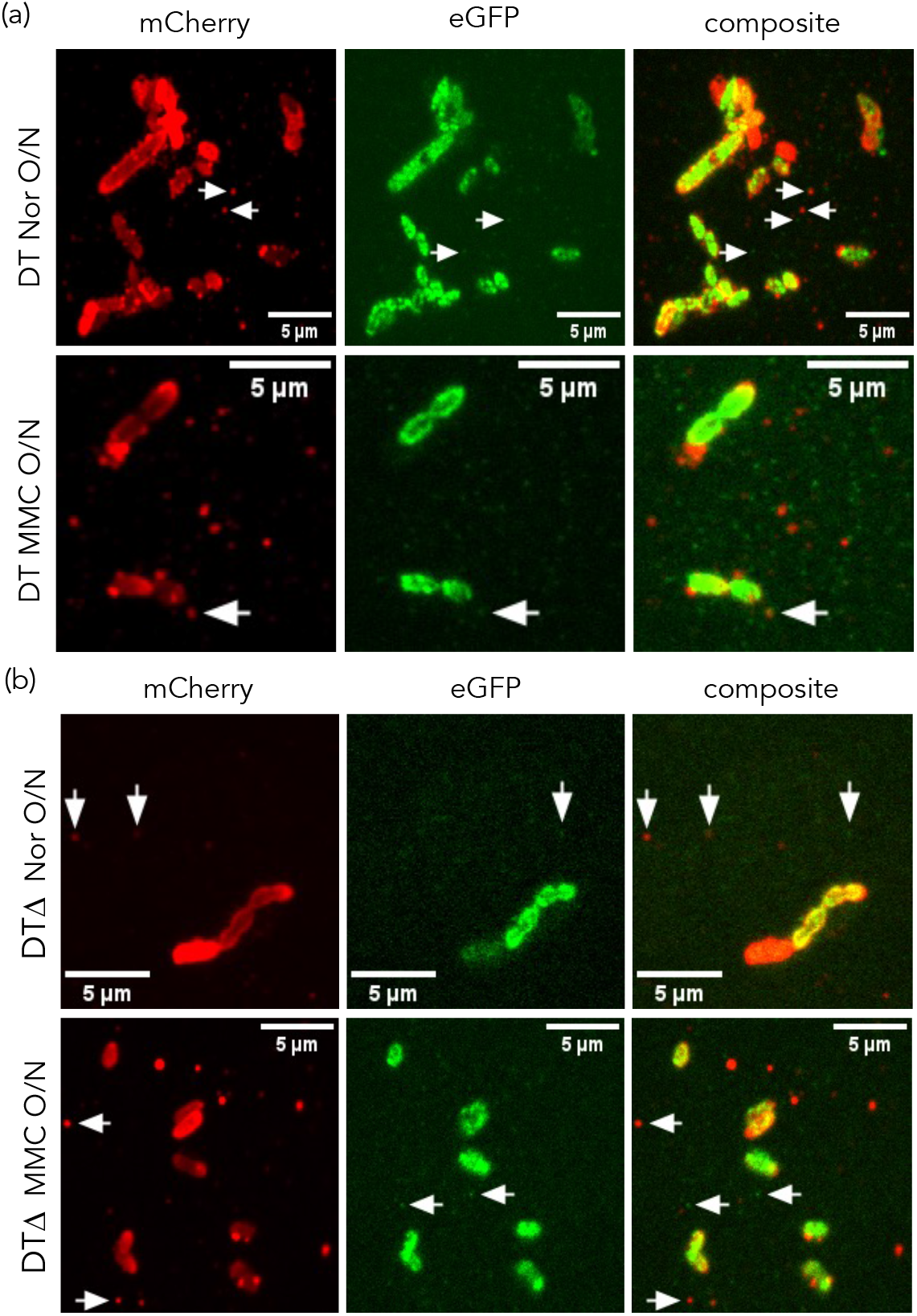
Effect of DNA damage-inducing compounds on membrane integrity after overnight exposure. (a) DT norfloxacin and mitomycin C-treated cultures. Arrows in the norfloxacin-treated sample show evidence of outer and inner membrane damage, and arrow in the mitomycin C-treated sample shows an OIMV. (b) DTΔ norfloxacin and mitomycin C-treated cultures. Arrows show evidence of outer and inner membrane damage. Nor – norfloxacin, MMC – mitomycin C, O/N – overnight.

**Figure 7.**
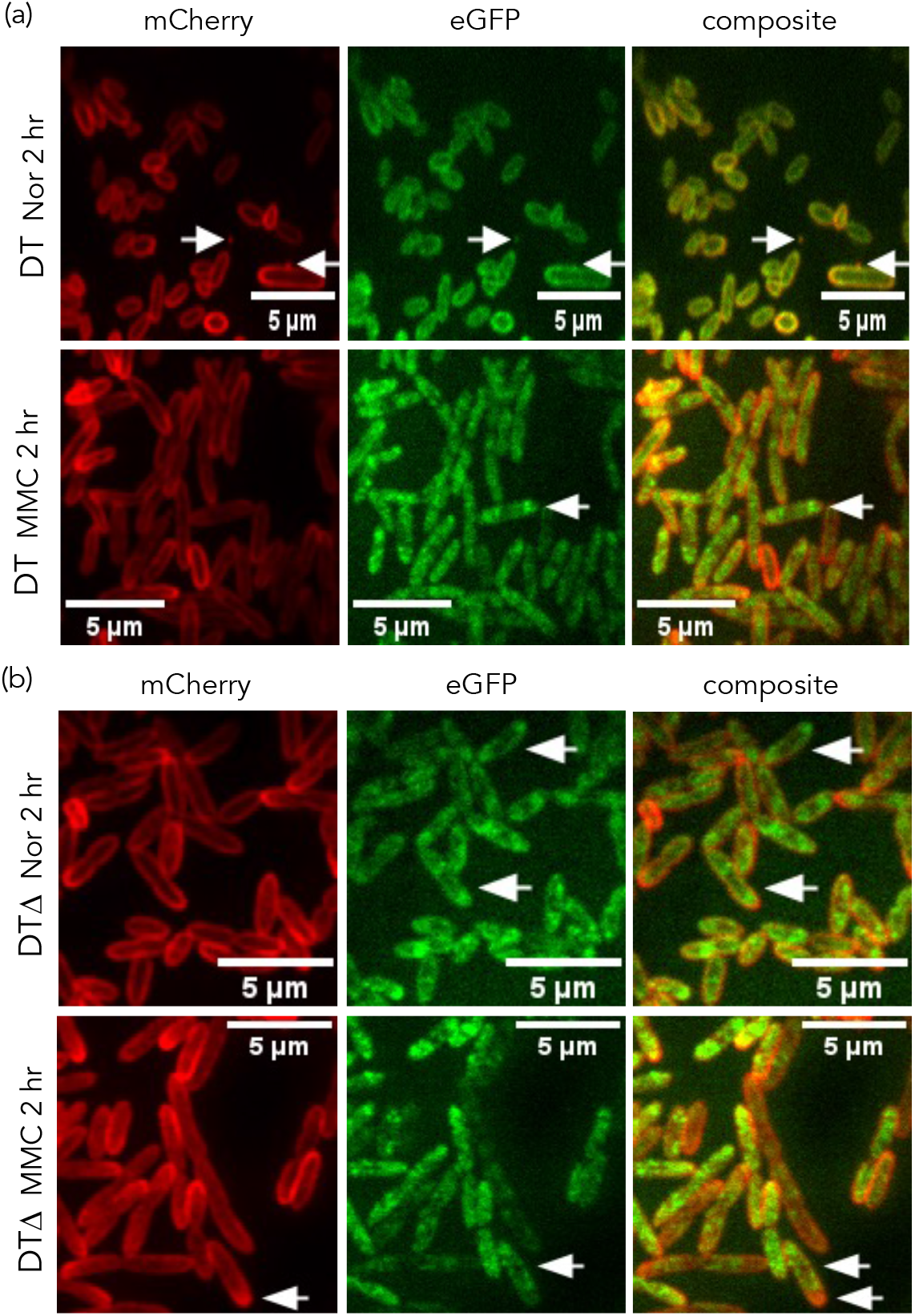
Effect of DNA damage-inducing compounds on membrane integrity after 2 hours of exposure. (a) DT norfloxacin and mitomycin C-treated cultures. Left arrow in the norfloxacin-treated sample shows an OIMV, while right arrow shows an OIMV in the process of being released. Arrows in the MMC-treated sample show polar re-localisation of inner membrane (AtpG) proteins. (b) DTΔ norfloxacin and mitomycin C-treated cultures. Arrows show polar re-localisation of outer membrane (Ax21) and inner membrane (AtpG) proteins. Nor – norfloxacin, MMC – mitomycin C.

TEM images show evidence of cell lysis in all fields of view for WT cultures treated with norfloxacin and mitomycin C – indications of cell lysis are also clearly observable in Δ*mal* samples, but not as much as the corresponding WT samples. Noticeably, the size of most MVs in both sets of samples is rather small (around 50 nm), which is in apparent contradiction to the data obtained from the SPT data; however, this was expected given the optimal size range of the instrument used for measurements. OIMVs were visible predominantly in WT samples, and these could be distinguished from other large MVs based on their staining profiles. Curiously, we also imaged some large single-layered vesicles that enclose some matter that stains similar to that observed in OIMVs – we speculate that these are cytoplasmic membrane vesicles (CMVs) (Figures 8–11, Supplementary Material 4).

**Figure 8.**
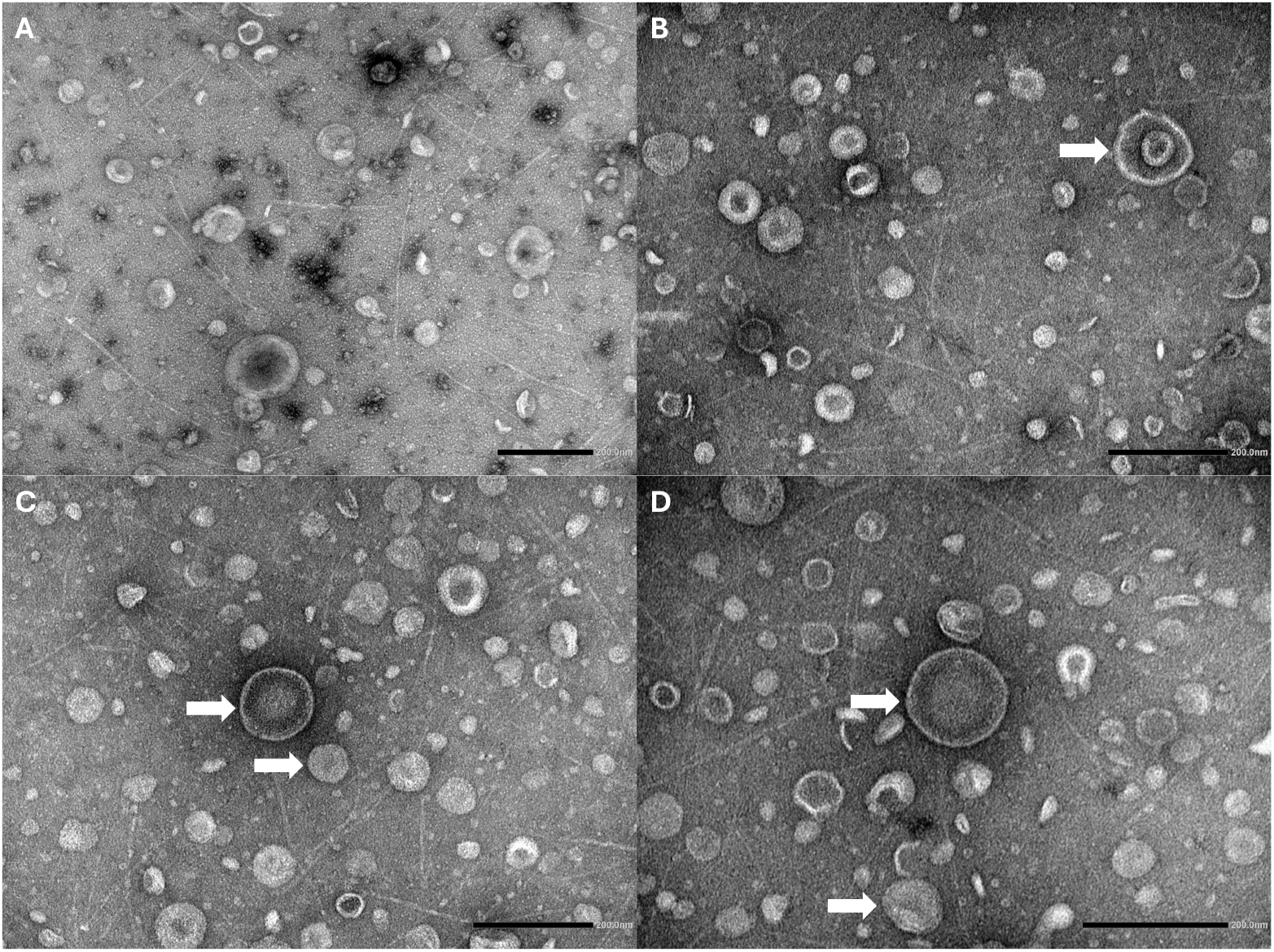
TEM images of WT cultures exposed to norfloxacin. (A) General field of view. Many phage tail-like particles are visible in the background. (B) An OIMV is clearly visualised (white arrow). (C) and (D) Two single-layered vesicles are visualised with different staining profiles – these correspond to CMVs (white arrows, above) and OMVs (white arrows, below). Scale bar – 200 nm.

**Figure 9.**
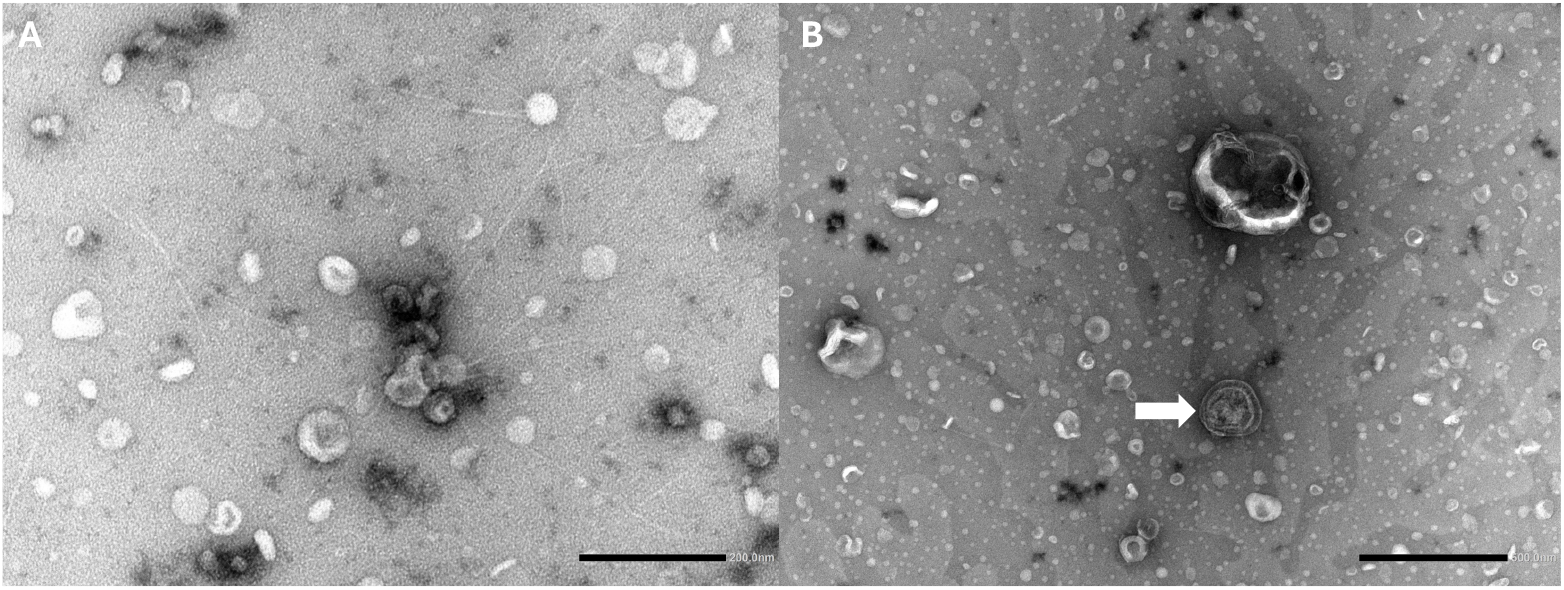
TEM images of WT cultures exposed to mitomycin C. (A) General field of view. Most vesicles are single-layered and contain lightly-stained matter. (B) An OIMV is clearly visible (white arrow). Scale bar – 200 nm.

**Figure 10.**
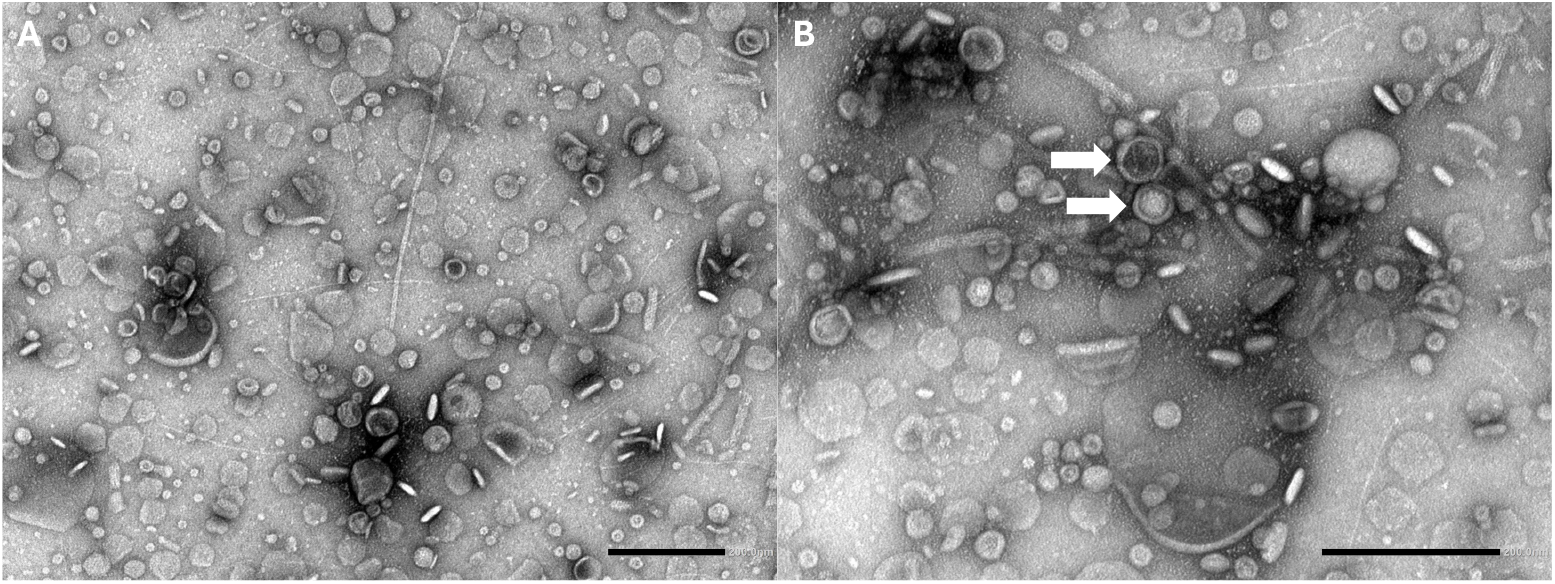
TEM images of Δ*mal* cultures exposed to norfloxacin. (A) General field of view. Many small particles are visible in addition to vesicles, likely corresponding to cell debris. (B) Two single-layered vesicles are visualised with different staining profiles – these correspond to a CMV (white arrow, above) and an OMV (white arrow, below). Scale bar – 200 nm.

**Figure 11.**
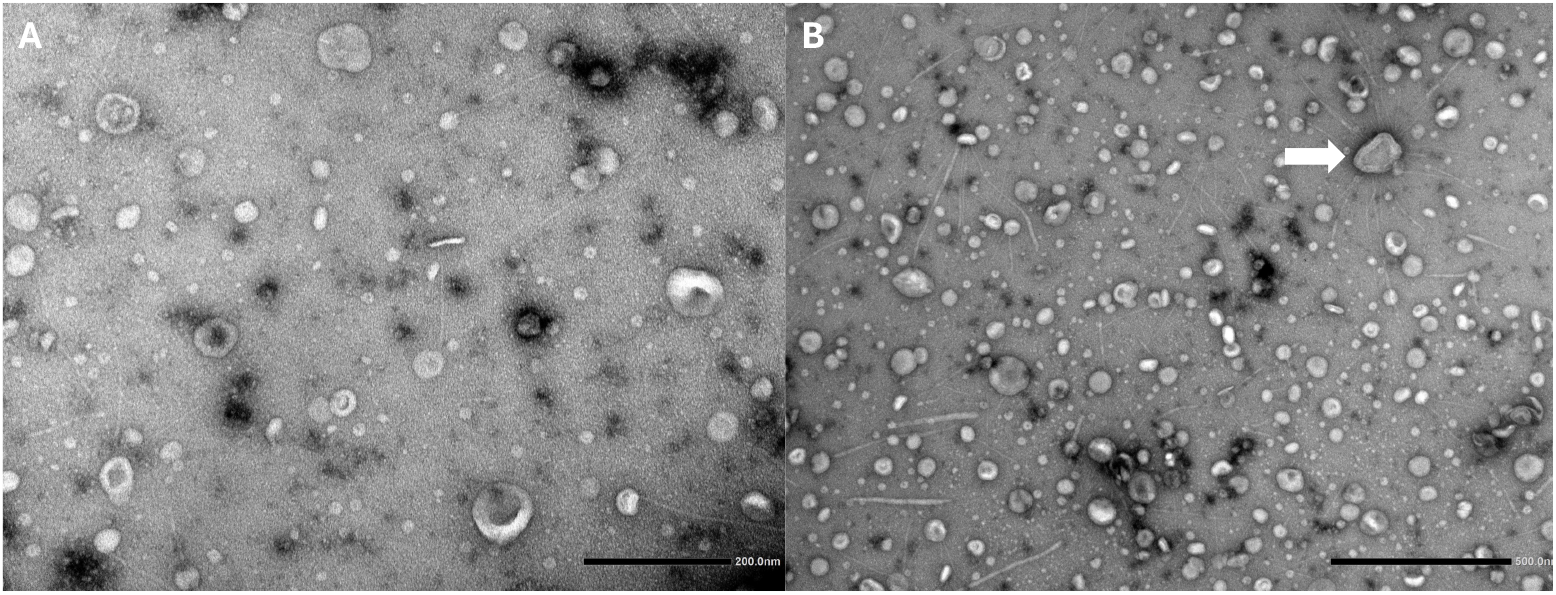
TEM images of Δ*mal* cultures exposed to mitomycin C. (A) General field of view. Larger vesicles are not common. (B) An OIMV with several phage tail-like particles attached to its surface is visible (white arrow). Scale bar – (A) 200 nm, (B) 500 nm.

Taking into account all our data, we posit that OMVs formed by self-assembly of shattered outer membrane fragments (i.e., E-OMVs) form the bulk of MVs produced by explosive cell lysis, not OIMVs – due to the inherent randomness of the nature of their biogenesis, E-OMVs of highly variable size and morphology are formed (Box 1). Furthermore, CMVs might also form in this manner, suggesting that cell lysis and concomitant re-circularisation of membrane fragments represents a pathway for the formation of all types of MVs in Gram-negative bacteria.

#### Box 1.

**Explanation for the abundance of E-OMVs**.

The surface area-to-volume (SA/V) ratio is an important determinant of cell size. As long as cells have enough surface area to support their growing volume, they are able to grow normally. When the surface area is unable to accommodate the increase in volume (i.e., when the SA/V reaches a critical minimum), the cell divides to produce daughter cells with a high SA/V ratio which facilitates their growth once again. This SA/V homeostasis is widely regulated by cells in all domains of life, such that their morphologies are suited for their respective environmental niches ^33^. Similarly, MVs are also subject to SA/V constraints – if the SA/V ratio is too low (i.e., if the MV is large), it is prone to rupturing under the tension of its own membrane or due to external forces.

The distance between the Gram-negative outer and inner membranes (OM and IM respectively) likely varies between bacteria (and even between cells). Let us assume that this distance in *S. maltophilia* is X nm. In the simplest scenario, this means that for an OIMV of a given size to exist, the OM must have a radius that is larger than the radius of the IM by approximately X nm. Furthermore, since the OM and IM are structurally linked by several mechanisms (including integral proteins that span both membranes like the Tol-Pal system), this periplasmic distance should remain roughly X nm in unchanging conditions. If the OM-IM distance increases too much (i.e., if the OM is much larger than the IM), the structural stability of the OIMV is compromised and it either collapses on itself or ruptures under the resulting tension. This is also influenced by the lipopolysaccharide (LPS) layer, an excess of which increases pressure towards the inside of the vesicle, and the peptidoglycan layer, to which certain proteins anchor and exert an inward-facing force. Instability is only exacerbated for smaller MVs, where surface forces dominate over bulk properties – this explains why stable OIMVs are larger than typical OMVs. To summarise, a stable OIMV needs to maintain an optimal periplasmic distance and have a lower SA/V ratio to reduce membrane tension. However, for an OMV of the same (or comparable) size, the only constraint is the SA/V ratio – since OMVs only have a single layer, they can stably exist across a much wider size range. Thus, the spontaneous re-assembly of membrane fragments upon cell lysis favours the production of E-OMVs. In additional support of this notion, it has also been proposed that unstable OIMVs can re-arrange to produce ‘second-generation’ E-OMVs ^3^.

It should be noted that these calculations are theoretical and do not factor in the inherent randomness associated with explosive cell lysis, wherein membrane fragments of unpredictable sizes and configurations are generated, nor do they account for the type of OIMVs previously reported that appear to resemble smaller CMVs contained within larger OMVs. Other studies have found heterogenous conformations of OIMVs where the OM-IM gap has been observed to be quite large; however, these could also be artefacts due to sample preparation and might not necessarily be representative of their structure under physiological conditions ^11^. Stable OIMVs are most likely the product of re-assembly of intact OM-IM fragments that satisfy the aforementioned criteria. OIMVs with large periplasmic spaces have been observed; however, their stability remains to be assessed ^34^.

Finally, we explored the spatiotemporal dynamics of LysSM in cells as a response to antibiotic-induced DNA damage – to this end, a standalone eGFP-tagged LysSM mutant and an mCherry-tagged Ax21 double mutant were constructed. A sub-MIC of norfloxacin was added to an exponentially-growing culture of the *ax21:mCherry mal:egfp* mutant, and the culture was allowed to grow for an additional 2 hours. Fluorescence microscopy showed little-to-no change in outer membrane stability (based on Ax21 localisation); however, LysSM was observed to be present as bright foci at the poles of most cells. At this time point, an excess of MVs was not noted in any of the fields of view imaged (Figure 12). Wanting to visualise the real-time upregulation of *mal*, we then subjected the *mal:egfp* mutant culture to a high concentration of norfloxacin (10 μg/ml) and imaged cells immediately after and up to a duration of 1 hour post-exposure using time-lapse microscopy. During the initial minutes, little signal was observed; however, after approximately 15 minutes, several transient but non-stochastic spots were visible in multiple cells, similar to those observed after extended growth in the presence of the lower concentration of norfloxacin. In most cases, multiple foci were observed per cell and they tended to appear most prominent at the poles (Supplementary Material 5). We hypothesise that these correspond to explosive lysis events, the products of which are a heterogenous mixture of OIMVs, E-OMVs, and possibly CMVs.

**Figure 12.**
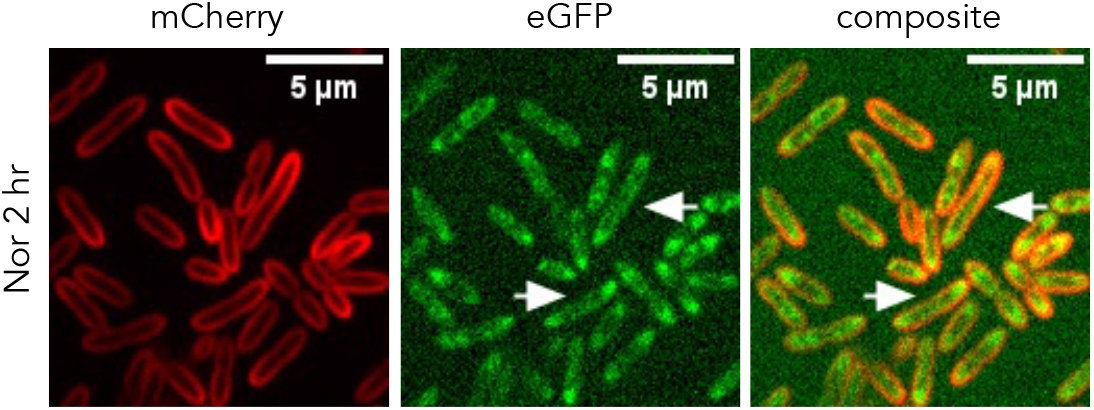
Effect of sub-MIC of norfloxacin on membrane integrity after 2 hours of exposure. The *ax21:mCherry mal:egfp* mutant was exposed to norfloxacin for 2 hours to observe the upregulation of *mal*. Arrows show polar localisation of LysSM. Nor – norfloxacin.

## Discussion

The discovery of MVs is relatively recent, and the elucidation of their biogenesis pathways even more so. While extensive information is available on the classical OMV, our understanding of other types remains limited. Antibiotic stress has been shown to be a major positive regulator of vesiculation in bacteria, with different classes stimulating the production of distinct types of MVs. Only recently was it demonstrated that exposure to DNA-damaging compounds was linked to the production of OIMVs, a novel type of MV characterised by the presence of both outer and inner membranes. The study showed that a latent bacteriophage in *P. aeruginosa* was induced in an SOS-dependent manner, leading to the production of a cell wall-degrading endolysin and consequent ‘explosive’ cell lysis, with the spontaneous re-assembly of membrane fragments producing OIMVs ^14^.

In prior research, it was observed that treatment of *S. maltophilia* with ciprofloxacin similarly resulted in the production of OIMVs, along with several phage tail-like particles (i.e., tailocins) ^15^. We assessed the role of the endolysin encoded by this cryptic prophage gene cluster (LysSM) in the context of membrane integrity and vesiculation capacity. A decrease in cell lysis was observed in a LysSM-deficient mutant relative to the WT, corresponding to improved growth and reduced biofilm formation, both of which are concurrent with expectations. However, the complete absence of cell lysis was not noted in the Δ*mal* mutant, and OIMVs were visible in these cultures as well, indicating that there exists some other trigger that is also possibly activated in a RecA-dependant manner. The pyocin cluster in *P. aeruginosa* was found to also code for a holin that was observed to increase cell lysis by translocation of the *lys* endolysin to the peptidoglycan layer ^14^. A definitively-annotated homologous holin was not found within the *S. maltophilia* K279a reference genome; however, holins form a diverse group of proteins, so this is not surprising ^35^. Genome mining yielded multiple candidates that might fulfil a similar role, including *smlt1851* (putative muramidase similar to that from the bacteriophage ASPE-1) and *smlt1944* (putative transmembrane muramidase similar to that from bacteriophage PS119), as well as several hypothetical proteins from the maltocin gene gluster (*smlt1039*–*smlt1065*). Further research is necessary to elucidate the roles of the genes that form these tailocin clusters.

A previous study in *P. aeruginosa* linked decreased MV production to various pyocin deletions ^36^. Following this, we demonstrated that not only is MV production in response to DNA damage impaired in the Δ*mal* mutant, but the size of MVs is also significantly reduced. The larger MVs secreted by the WT were previously thought to predominantly be OIMVs; however, using fluorescence microscopy, we observed little overlap between signals corresponding to outer and inner membrane fragments. As a result, we concluded that most MVs produced through LysSM-mediated explosive cell lysis are E-OMVs, a distinct type of OMV formed by the re-arrangement of outer membrane fragments. Data from our fluorescence microscopy experiments appear to suggest that the re-arrangement of inner membrane fragments also occurs, resulting in the production of CMVs – this is also corroborated by data from TEM images that show single-layered vesicles with distinct staining profiles. CMVs have been reported in Gram-negative bacteria but their biogenesis remains unclear ^3^. It is not possible to provide a definitive answer about the presence of CMVs as yet, since sub-populations of MVs can neither be easily distinguished or separated using current technology. Using metal nanoparticles that are specific to each membrane in conjunction with TEM might be one way to distinguish MV types, although this would not be entirely quantitative ^37^. Flow cytometry might be another possible method to do so, as demonstrated by a recent study ^38^. However, current hardware limitations make separation of nanometre scale particles very cumbersome, necessitating further developments in the field before they can be routinely utilised in the MV biology.

Another finding we (inadvertently) reported was the change in localisation of membrane proteins in response to antibiotic challenge. The selected outer and inner membrane proteins (Ax21 and AtpG respectively) appeared to distribute themselves to the poles when exposed to cell wall-targeting and DNA-damaging compounds. While it is possible that this re-localisation is purely coincidental and selection of different membrane proteins would have provided different results, it is nonetheless worth examining the roles of these proteins. Structurally, the mature form of Ax21 is a typical porin composed of a transmembrane β-barrel with 10 antiparallel strands – given that it localises in the outer membrane, it is likely present as a homotrimer. Porins represent a diverse group of molecules involved in, among other things, the passive transport of hydrophilic molecules (including antibiotics such as β-lactams and fluoroquinolones) ^39^. Due to the roughly cylindrical architecture of bacilli, the outer membrane is less fluid towards the poles (where the curvature is more). Lateral diffusion of porins within the membrane can cause their accumulation in these regions, thereby restricting uptake of these molecules ^40^. Furthermore, if MVs are preferentially produced at the poles, this provides cells an alternate way of membrane remodelling by shedding these porins on the surface of OMVs. The exact role of Ax21 in *S. maltophilia* is controversial due to a recent paper retraction ^41^. Our observation provides some insight into its function in response to antibiotic stress. On the other hand, AtpG is a well-characterised component of the bacterial ATP synthase complex ^42^. Due to its involvement in ATP synthesis, polar localisation in DNA damage-associated cells might be seen as a cellular response to meet energy demands in nascent daughter cells ^43^. We speculate that this could represent a sort of ‘jump ship’ mechanism of cell division that de-prioritises repairing damaged DNA, wherein a single healthy daughter cell is produced while the other daughter cell harbours the damaged parental DNA. In support of this is the fact that exposure to cell wall-targeting antibiotics (which suppress cell division) causes AtpG to localise instead as foci throughout the cell. Research into such a mechanism could be an interesting area for future studies and antibiotic development.

Finally, the upregulation of latent bacteriophages could prove useful as a therapeutic method for the treatment of stubborn infections. *S. maltophilia* MVs produced in response to DNA damage have been observed to be enriched for tailocins which are associated with their production ^15^. These particles have been shown to exert antibacterial activity against multiple Gram-negative and Gram-positive species ^18^. Given that exposure to only a sub-MIC of antibiotic is necessary to stimulate their production, one could imagine a counter-intuitive strategy wherein small amounts of DNA-damaging compounds are administered not to kill the bacteria themselves, but rather to cause the release of tailocins that kill other bacteria present in the same niche. This could be particularly useful in the context of polymicrobial infections, such as those associated with cystic fibrosis. Since MVs have also been shown to be major components of bacterial biofilms, stimulation of DNA damage-associated tailocin-enriched MVs within these structures might present itself as an interesting course of treatment, perhaps as an extension of current phage therapy ^44,45^.

## Data availability statement

Optimised protocols for all experiments and additional microscopy images can be found within the manuscript and the corresponding Addenda. Further data can be provided by request to the corresponding author.

## Author contributions

DC conceptualised and carried out all experiments (unless otherwise stated) and wrote the manuscript. JV partly performed membrane vesicle-related experiments and contributed to writing of the manuscript. SS, KR, and KB contributed to membrane vesicle quantification experiments. BD supervised the work, assisted in experiment conceptualisation, and reviewed the manuscript.

## Acknowledgments

The authors express their gratitude to the Ghent University Light Microscopy (GLiM) Core at Ghent University for the use and support on the spinning disk microscope, and Femke Baeke from the Vlaams Instituut voor Biotechnologie (VIB) BioImaging Core (Ghent) for her technical support with the electron microscope.

## Ethical statements and declarations

Funding was obtained from the Fonds Wetenschappelijk Onderzoek (FWO) grant number G005719N, the Belgian Federal Science Policy Office (Belspo), and the European Space Agency Programme de Développment d’Expériences scientifiques (ESA PRODEX) as part of the Yeast Nanomotion Project. This manuscript does not contain studies with human participants or animals performed by any of the authors.

