## Supplementary Material 1-4 for "DNA damage-associated vesicle production in *Stenotrophomonas maltophilia* is mediated by a cryptic tailocin endolysin"

**Supplementary Material 1. List of screening primers used.**

| **Type of primer** | **Primer sequence (5’ 🡪 3’)** |
| --- | --- |
| *egfp* gene-specific | FP: CACATGAAGCAGCACGACTT  RP: TGCTCAGGTAGTGGTTGTCG |
| *atpG* gene-specific | FP: TGTGCTTTGAACGAACGCGG  RP:TGATTTCGCGTCCGCTTGCCAT |
| *mal* gene-specific | FP: CATTGATGGTGATGCCGCG  RP: AGACGCTCTGTTGAACCTGG |
| *mal* flanking region | FP: CATTGATGGTGATGCCGCG  RP: AGACGCTCTGTTGAACCTGG |
| *ax21* gene-specific | FP: ATGAAGAATTCGCTGATTGCTCTGG  RP: GCGGAATCCAATTCGTTGCAAT |

**Supplementary Material 2. Sequences of homology regions and inserts used.**

**Δ*mal* and *mal:egfp* (1107786–1108271, minus-strand)**

Upstream homology sequence (1107585–1107785, plus-strand)

CGCGCCTGCGGCGCTGTTGGCGAGGGCCTGCTGGTGCGCGCGCTGGACTGCATCGGTGCGGGCCTCCTGCTGCGCTGCCCGGGTGGTGGCAACGGCAAGGTCCGCAGTGCGGTCGCGCAGGGTCCATCCCAGCCAGGCGCAGCTCGCGTGGCTACCGGCAAGGACCAGCAACCCGATGCCCAACCTCAGTCCCGCGGGCG

Downstream homology sequence (1107788–1108088, plus-strand)

CTGCAGGGCTCCGCCGGCGGCACGGTACGCCGCGCGCAGTGTTTCCAGTGCGTGCTCCTTCTGGCCGTAGCCGGCGCCCGGCAGCGATGCCCAGATGCGCCGGGCCGCGGTGACGGCGGCATCGAAGCGTCCCAGCCGGATCAGCTCGTAGGCACCGCACTGTTTCAGCAGGGCGACTGCTGCACGGTCCTGGGAGACTGGCCCGAAGTCAGGCAGACCCAGTCGCGCGCGCAGGTCATCCCAGGTGCTGCGCAGGAACTGGTAGCGGCCGGCGGCGCTGGATTTGATGCCATAGCGTGG

Linker + eGFP sequence (reverse complement)

TTACTTGTACAGCTCGTCCATGCCGAGAGTGATCCCGGCGGCGGTCACGAACTCCAGCAGGACCATGTGATCGCGCTTCTCGTTGGGGTCTTTGCTCAGGGCGGACTGGGTGCTCAGGTAGTGGTTGTCGGGCAGCAGCACGGGGCCGTCGCCGATGGGGGTGTTCTGCTGGTAGTGGTCGGCGAGCTGCACGCTGCCGTCCTCGATGTTGTGGCGGATCTTGAAGTTCACCTTGATGCCGTTCTTCTGCTTGTCGGCCATGATATAGACGTTGTGGCTGTTGTAGTTGTACTCCAGCTTGTGCCCCAGGATGTTGCCGTCCTCCTTGAAGTCGATGCCCTTCAGCTCGATGCGGTTCACCAGGGTGTCGCCCTCGAACTTCACCTCGGCGCGGGTCTTGTAGTTGCCGTCGTCCTTGAAGAAGATGGTGCGCTCCTGGACGTAGCCTTCGGGCATGGCGGACTTGAAGAAGTCGTGCTGCTTCATGTGGTCGGGGTAGCGGCTGAAGCACTGCACGCCGTAGGTCAGGGTGGTCACGAGGGTGGGCCAGGGCACGGGCAGCTTGCCGGTGGTGCAGATGAACTTCAGGGTCAGCTTGCCGTAGGTGGCATCGCCCTCGCCCTCGCCGGACACGCTGAACTTGTGGCCGTTTACGTCGCCGTCCAGCTCGACCAGGATGGGCACCACCCCGGTGAACAGCTCCTCGCCCTTGCTCACCATGCTGCCGCTGCCGCTGCC

***atpG:egfp* (4219920–4220783, minus-strand)**

Upstream homology sequence (4219080–4219919, plus-strand)

ACGGCATGCGGCCCAGCAGTGCCGACACTTCGGTACCGGCCAGGGTGTAGCGGTAGATGTTGTCGACGAACAGCAGGACGTCCTTGCCCTTGCCGTTTTCGTCCTTCTCGTCGCGGAAGTACTCGGCCATGGTCAGGCCGGTCAGGGCGACGCGCAGACGGTTGCCCGGCGGCTCGTTCATCTGGCCGTACACCATCGCCACCTTGTCGAGGACGTTGGAGTCCTTCATTTCGTGGTAGAAGTCGTTGCCCTCACGGGTACGCTCACCCACGCCGGCGAACACGGACAGACCGCTGTGCGCCTTGGCGATGTTGTTGATCAGCTCCATCATGTTGACGGTCTTGCCGACGCCGGCGCCGCCGAACAGGCCGACCTTGCCGCCCTTGGCGAACGGGCACATCAGGTCGATGACCTTGATGCCGGTTTCCAGCAGTTCGGTGGCCGGGGACTGGTCTTCGTACGACGGGGCCGCACGGTGGATTTCCCAGCTGTCGCTGGCGGCCACCGGGCCGGCTTCGTCGATCGGACGGCCGAGCACGTCCATGATGCGGCCCAGGGTGCCGGCGCCGACCGGCACCGAGATGCCACGGCCGGTGTTGACGGCAACCAGGTTGCGCTTCAGGCCGTCGGTGGAACCGAGGGCGATGGTACGCACCACGCCGTCGCCCAGCTGCTGCTGGACTTCGAGGGTGATCTCGGTGTTTTCCACCTTCAGTGCGTCGTACACCTTCGGCACCGATTCACGCGGGAATTCGACGTCGACGACCGCGCCGATGATCTGAACGATCTTGCCCTGACTCATTGCTGCATCCTCTAATGTGTGCTTTGAACGAACGCGG

Downstream homology sequence (4219917–4220783, plus-strand)

GACTGCTGCCGCGCCGCCGACGATTTCGGAGATTTCCTGGGTGATCGCTGCCTGGCGCGCCTTGTTGTAGACGAGCTGCAGGGTGCCGATCAGCTTGTTGGCGTTGTCGCTCGCCGCCTTCATCGCAACCATGCGTGCCGCATGCTCGGAGGCGACGTTTTCCAGCAGTGCCTGGTACACCAGCGACTCGATGTAACGCGTCATCACGTGCTCGAGCACGGTCGCGGCATCGGGTTCGTACAGGTAATCCCAGTCGTGGTGAGCGACCTGCTTCTCAGCCGGCGGCAGCGGCAGCAGCTGGTCGAAGCTGGCCTTCTGCACCATGGTGTTCACGAAGCGGTTGTAGACCAGGTACACGCGGTCGATCTTGCCTTCGGTGAAGGCGTCGAGCATGACCTTGATCACACCGATCAGCGATTCCAGCTTCGGCACATCGCCGATGTGGGTCACGCTGCCGACCATGTTGACCTTCACGCGGCGGAAGAAGGTCGAAGCCTTCTGGCCGATGGTCACCAGGTCCACTTCCGCACCCTTGTCCTGCCATGCCTTGGCTTCGCCCAGCATCTTGCGGAACAGGTTGTTGTTCAGGCCGCCGGCCAGGCCGCGATCGGAGGAGATCACGATGAAGCCGACCCGCTTGACCTGCTCGCGCTCGACCAGGAACGGATGCTGGTAGTCGGTGCTGGCCTGGGCCAGGTGACCGATCACCTGCTTCATCGCCTGCGCGTACGGACGCGAGGTCTTCATCCGATCCTGCGCCTTGCGGATCTTGGAGGCCGAGACCATTTCCAGGGCGCGCGTCACCTTGCGGGTGTTCTGCACGCTCTTGATCTTGGTTTTGATTTCGCGTCCGCTTGCCAT

Linker + eGFP sequence (reverse complement)

TTACTTGTACAGCTCGTCCATGCCGAGAGTGATCCCGGCGGCGGTCACGAACTCCAGCAGGACCATGTGATCGCGCTTCTCGTTGGGGTCTTTGCTCAGGGCGGACTGGGTGCTCAGGTAGTGGTTGTCGGGCAGCAGCACGGGGCCGTCGCCGATGGGGGTGTTCTGCTGGTAGTGGTCGGCGAGCTGCACGCTGCCGTCCTCGATGTTGTGGCGGATCTTGAAGTTCACCTTGATGCCGTTCTTCTGCTTGTCGGCCATGATATAGACGTTGTGGCTGTTGTAGTTGTACTCCAGCTTGTGCCCCAGGATGTTGCCGTCCTCCTTGAAGTCGATGCCCTTCAGCTCGATGCGGTTCACCAGGGTGTCGCCCTCGAACTTCACCTCGGCGCGGGTCTTGTAGTTGCCGTCGTCCTTGAAGAAGATGGTGCGCTCCTGGACGTAGCCTTCGGGCATGGCGGACTTGAAGAAGTCGTGCTGCTTCATGTGGTCGGGGTAGCGGCTGAAGCACTGCACGCCGTAGGTCAGGGTGGTCACGAGGGTGGGCCAGGGCACGGGCAGCTTGCCGGTGGTGCAGATGAACTTCAGGGTCAGCTTGCCGTAGGTGGCATCGCCCTCGCCCTCGCCGGACACGCTGAACTTGTGGCCGTTTACGTCGCCGTCCAGCTCGACCAGGATGGGCACCACCCCGGTGAACAGCTCCTCGCCCTTGCTCACCATGCTGCCGCTGCCGCTGCC

***ax21:mCherry* (384354–384926, plus-strand)**

Upstream homology sequence (384021–384412, plus-strand)

CACGCGTTCGGCGCGCTGGTAGTCGGCCGCCTGCAGCGCGTGTGCGGTGTGTGGCGACAAGAGGGCAACGCATGCAAAGACAGCGGCGCGGACACACGTCCGCAGCGGATCGGAAACGAGCAAGGCAACCTCCTGTTGCAATCGTCGAAGGATTACAGATGCACGCGCGCCGCGCGCGTATGCCAGACGGAGCCTGAATCAAGCCCAAAAATTCAGCAATGCGAAGACCACTGTTTTCAGGGAGCGTTGTGGCGAATAACGGAGAATTTACATTCCCGATGCGATGGCGCCCTGGCGCTGCTGCACACCATAAGAAAGTAAAGGTGTACCCCCATGAAGAATTCGCTGATTGCTCTGGCCCTGGCCGCTGCCCTGCCGTTCACCGCTTCGGCT

Downstream homology sequence (384927–385427, plus-strand)

TTGCTTCCTCGGCGCGCCTGCGCGCTGAAGATGCCTCGACAGGCCCGGCCCATGCCGGGCCTGTTGCGTTTCGGGCTGGGCAAACGCTACCTGACCGTGTTGAGGTGGACGGGCGGCATTGCAACGAATTGGATTCCGCAGGCGTCGGCTGCGCCCTAGGGTGAGTGGCCTGTCCTGCAGGAATGCAACGATGACCCCGTTTCCCTCGCCCCGCGATTCTCCCTGCCGCTTGTTGCTGCTGGATCCGCATCCACTGTTGCGGCATGGCGTGCAGGAACTGCTGGCCCGGCAGGCGGGCCTGCAGATCGATGGAAGCTATGGCCGCAGCGGTGATCTGCTGCAGCGCCTGCAGCAGCAGGCAGCTGCGATCGATCTGCTGCTGGTTGACGCCTTGCCGATCGGTGACAGCGGGGAAGGCCTGGAACTGCTGCGGCACGTGTCACGACATTGGCCGGCAGTGCCGATCCTGGTGCTGTCCGCACACTGCAACGCGGGTGTCG

Replacement gene (mCherry sequence + linker + smlt0387)

ATGAAGAATTCGCTGATTGCTCTGGCCCTGGCCGCTGCCCTGCCGTTCACCGCTTCGGCTATGGTGAGCAAGGGCGAGGAGGATAACATGGCCATCATCAAGGAGTTCATGCGCTTCAAGGTGCACATGGAGGGCTCCGTGAACGGCCACGAGTTCGAGATCGAGGGCGAGGGCGAGGGCCGCCCCTACGAGGGCACCCAGACCGCCAAGCTGAAGGTGACCAAGGGTGGCCCCCTGCCCTTCGCCTGGGACATCCTGTCCCCTCAGTTCATGTACGGCTCCAAGGCCTACGTGAAGCACCCCGCCGACATCCCCGACTACTTGAAGCTGTCCTTCCCCGAGGGCTTCAAGTGGGAGCGCGTGATGAACTTCGAGGACGGCGGCGTGGTGACCGTGACCCAGGACTCCTCCCTGCAGGACGGCGAGTTCATCTACAAGGTGAAGCTGCGCGGCACCAACTTCCCCTCCGACGGCCCCGTAATGCAGAAGAAGACCATGGGCTGGGAGGCCTCCTCCGAGCGGATGTACCCCGAGGACGGCGCCCTGAAGGGCGAGATCAAGCAGAGGCTGAAGCTGAAGGACGGCGGCCACTACGACGCTGAGGTCAAGACCACCTACAAGGCCAAGAAGCCCGTGCAGCTGCCCGGCGCCTACAACGTCAACATCAAGTTGGACATCACCTCCCACAACGAGGACTACACCATCGTGGAACAGTACGAACGCGCCGAGGGCCGCCACTCCACCGGCGGCATGGACGAGCTGTACAAGGGCAGCGGCAGCGGCAGCGCTGAGAACCTGTCGTACAACTACGCTGAAGCCGACTACGCCAAGACCGACGTCGATGGCATCAAGGCTGACGGCTGGGGCGTCAAGGGTTCCTACGGCTTCCTGCCGAACTTCCACGCGTTCGGTGAATACAGCCGTCAGGAAGTCGACCACACCAACATCAAGGTTGACCAGTGGAAGGTCGGTGCCGGCTACAACGTCGAAATCGCTCCGTCGACCGACTTCGTTGCCCGCGTTGCCTACCAGAAGTTCGATCGCAAGCACGGCCTGGACTTCAACGGCTACAGCGCTGAAGCCGGTATCCGCACCGCCTTCGGTGCCCACGCCGAGGTCTACGGCATGGTCGGCTACGAAGACTACGCCAAGAAGCACGGCGTCGACATCGACGGCCAGTGGTACGGCCGCCTGGGTGGTCAGGTCAAGCTGAACCAGAACTGGGGCCTGAACGGCGAGCTGAAGATGAACCGCCACGGCGACAAGGAATACACCGTCGGCCCGCGCTTCAGCTGGTAA

**Note:** All genomic coordinates are with reference to the genome of strain K279a – partial genomic data for strain 44/98 is available, but not published.

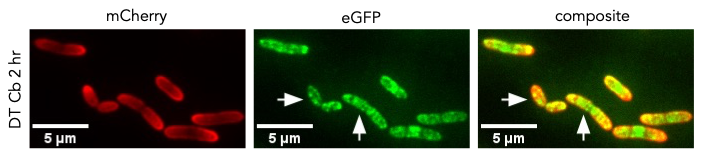
**Supplementary Material 3. Additional fluorescence microscopy images.**

**Figure 1. Effect of carbenicillin on membrane integrity after 2 hours of exposure.** Images of the DT carbenicillin-treated culture. Arrows show foci at which AtpG (inner membrane) is concentrated.

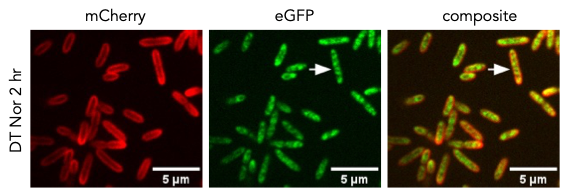
Cb – carbenicillin.

**Figure 2. Effect of norfloxacin on membrane integrity after 2 hours of exposure.** Images of the DT norfloxacin-treated culture. Arrows show foci at which AtpG (inner membrane) is concentrated.

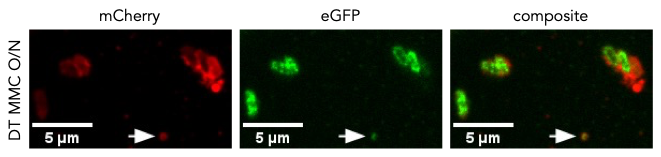
Nor – norfloxacin.

**Figure 3. Effect of mitomycin C on membrane integrity after overnight exposure.** Images of the DT mitomycin C-treated culture. The arrow shows a large OIMV with co-localised mCherry (outer membrane) and eGFP (inner membrane) signals.

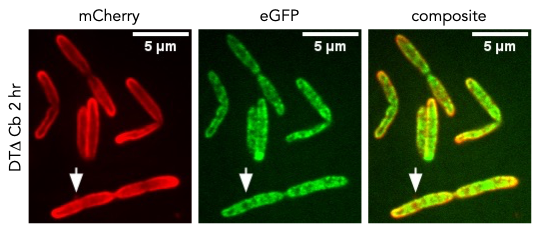
MMC – mitomycin C.

**Figure 4. Effect of carbenicillin on membrane integrity after 2 hours of exposure.** Images of the DTΔ carbenicillin-treated culture. Arrows shows an OMV with only mCherry (outer membrane) signal.

Cb – carbenicillin.

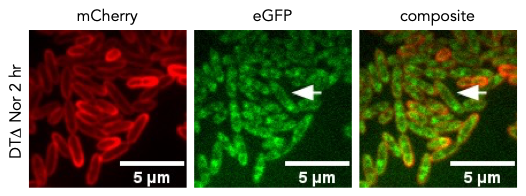
**Figure 5. Effect of norfloxacin on membrane integrity after 2 hours of exposure.** Images of the DTΔ norfloxacin-treated culture. Arrows show foci at which AtpG (inner membrane) is concentrated.

Nor – norfloxacin.

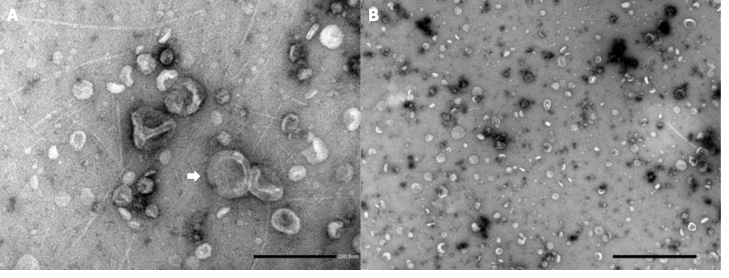
**Supplementary Material 4. Additional TEM images.**

**Figure 1. TEM images of *S*. *maltophilia* cultures exposed to DNA-damaging agents.** (A) An OIMV imaged in a WT culture exposed to norfloxacin (white arrow). (B) General field of view of a Δ*mal* culture exposed to mitomycin C. Most vesicles visible are small (50–100 nm) while only a few are large (more than 100 nm). Phage tail-like particles are visible in the background of both images.

Scale bar – (A) 200 nm, (B) 500 nm.
